# Genome-Wide CRISPR/Cas9 Screening Reveals Lipid Metabolism and Inflammatory Signalling as Modulators of Ganoderic Acid DM Cytotoxicity

**DOI:** 10.1101/2025.10.14.682027

**Authors:** Nimo Abdullah, Jacqueline Lewis, Prakash Arumugam

## Abstract

Ganoderic Acid DM (GA-DM), a triterpenoid derived from *Ganoderma lucidum*, exhibits anti-cancer and anti-diabetic activities, but the underlying mechanisms of action remain unclear. To identify genetic modulators of GA-DM response, we conducted a genome-wide CRISPR/Cas9 knockout screen in human melanoma cells. The screen revealed key roles for genes regulating lipid metabolism and inflammatory signalling, particularly the SREBP (Sterol Regulatory Element-binding Protein) and NF-κB (Nuclear Factor kappa-light-chain-enhancer of activated B cells) pathways, in the cellular response to GA-DM. While loss of genes involved in regulation of cholesterol biosynthesis conferred resistance to GA-DM, the disruption of ubiquitin-mediated proteolysis and the Hippo pathway genes sensitised cells to GA-DM. Inflammatory genes enriched at later time points suggests that a delayed cellular response contributes to cytotoxicity. Our findings propose a mechanistic model wherein GA-DM perturbs lipid and inflammatory pathways to exert cytotoxic effects and highlights potential targets to enhance its therapeutic efficacy. This work demonstrates the utility of functional genomics in elucidating natural product mechanisms and guiding rational drug development.

## Introduction

Mushrooms have been universally accepted as both food and medicine for centuries, with *Ganoderma lucidum*, commonly known as Reishi or Lingzhi, standing out as one of the most significant species in East Asian medicine (Ahmad, 2018; J. Liu et al., 2023). This mushroom has traditionally been associated with promoting longevity and enhancing vitality (Jong & Birmingham, 1992). In recent years, G. lucidum has attracted growing scientific interest due to its diverse pharmacological properties, largely attributed primarily to its bioactive triterpenoids and polysaccharides (Benzie IFF, Wachtel-Galor S et al., 2011; P.-G. Cheng et al., 2013).

Among the triterpenoids, Ganoderic Acids (GAs), particularly types A, B, C, D, H, R and Me, demonstrated a wide range of therapeutic effects (Liang et al., 2019). These lanostane-type compounds are known for their anticancer activity and have shown to induce apoptosis in cancer cells from breast, prostate, liver and lung origins (Müller et al., 2006; Sliva et al., 2002; Stanley et al., 2005; Tang et al., 2006; Wang et al., 2017; Xu et al., 2010). GAs can inhibit angiogenesis, metastasis and tumour invasion, by modulating key signalling pathways such as NF-κB, MAPK (mitogen-activated protein kinase) and PI3K (Phosphoinositide 3-kinase)/Akt(Ak strain transforming), all of which are critical for cancer cell survival and proliferation (S. Cheng & Sliva, 2015; Guo et al., 2020). However, the underlying mechanisms of these effects are complex and multifactorial. Some studies have reported that GAs can suppress the NF-κB pathway, which contributes to reduced inflammation and tumour progression and inhibit the PI3K/Akt and MAPK pathways, thereby blocking survival signals in cancer cells (Jiang et al., 2008). Another proposed mechanism involved the GA-mediated activation of the mitochondrial apoptotic pathway by triggering cytochrome c release and caspase activation (R. M. Liu & Zhong, 2011; Tang et al., 2006). Moreover, GAs were shown to inhibit topoisomerases, thereby disrupting DNA replication in rapidly dividing cells (C. H. Li et al., 2005). Beyond their anticancer properties, GAs have also demonstrated antiviral potential *in vitro*, particularly against hepatitis B virus (HBV) and human immunodeficiency virus (HIV) (Ahmad et al., 2024; Kang et al., 2015; Y. Q. Li & Wang, 2006; J. Liu et al., 2004).

Despite the therapeutic potential, GAs present several challenges for clinical application. Their poor water solubility and limited bioavailability hinder their efficacy *in vivo*, prompting research into delivery systems such as nanoformulations and liposomes to enhance pharmacokinetic performance (Karimi et al., 2022; Zhang et al., 2019). However, clinical studies investigating purified GAs remain limited. Most available human trials use *G. lucidum* extracts, which contain a mixture of triterpenoids, polysaccharides and other compounds. This makes it difficult to attribute the observed effects to GAs alone. For example, while one study showed an improved immune response in lung cancer patients undergoing chemotherapy via supplementation with *G. lucidum* extracts, other studies have reported variable outcomes on fatigue and quality of life in cancer patients (Gao et al., 2003; S. Wu et al., 2024; Zhao et al., 2012). Nevertheless, these findings are confounded by the lack of standardisation and the use of complex mixtures rather than isolated compounds.

Among the individual GAs, Ganoderic Acid A (GA-A) and Ganoderic Acid DM (GA-DM) have been especially well-studied. GA-A has demonstrated anti-hyperlipidemic properties by suppressing the expression of sterol regulatory element-binding proteins (SREBPs), which are key transcription factors involved in lipid biosynthesis (Zhu et al., 2018). Through this mechanism, GA-A reduced intracellular cholesterol and fatty acid levels *in vitro*. Additionally, GA-DM demonstrated significant anticancer potential, particularly in breast, melanoma and prostate cancer models (Hossain et al., 2012; Ruan et al., 2015; G.-S. Wu et al., 2012). For example, GA-DM has been shown to induce cytotoxicity in both androgen-dependent and androgen-independent prostate cancer cells (Liu et al., 2009). This effect was attributed to its inhibition of 5-α-reductase, the enzyme responsible for converting testosterone into the more potent androgen dihydrotestosterone (DHT). The structural similarity between GA-DM and DHT is thought to enable GA-DM to bind competitively to the active site of 5-α-reductase. By disrupting androgen biosynthesis and downstream signalling pathways, GA-DM may inhibit the proliferation and survival of androgen-responsive prostate cancer cells, contributing to its cytotoxic activity thus mirroring the effect of other steroidal inhibitors. Besides the effect of GA-DM on 5-α-reductase and androgen receptors, GA-DM was shown to inhibit osteoclastogenesis thereby limiting bone metastasis in prostate cancer (Liu et al., 2009).

Understanding the molecular mechanisms of these bioactive compounds is essential for optimising their therapeutic potential and guiding drug development. Functional genomic approaches, such as pooled CRISPR/Cas9 knockout screens, are valuable tools for identifying novel molecular targets. These screens use guide RNAs to direct the Cas9 endonuclease to specific DNA sequences, generating double-strand breaks and loss-of-function mutations through frameshift indels (Cong et al., 2013; Jinek et al., 2012; Mali et al., 2013). In this study, we conducted a genome-wide CRISPR/Cas9 knockout screen in human melanoma cells to identify genes that confer resistance or sensitivity to GA-DM. Leveraging the high-coverage Brunello library, we uncovered key components of lipid metabolism and inflammatory signalling, particularly the SREBP and NF-κB pathways, as modulators of GA-DM-induced cytotoxicity. These findings provide novel insights into the molecular mechanisms of GA-DM action, suggesting that its anticancer effects may be driven by disruption of cholesterol homeostasis and activation of inflammatory responses.

## Material and methods

The genome-wide CRISPR/Cas9 screen was performed according to the methods described previously (Goh et al., 2021).

### Plasmid library amplification

The Brunello human CRISPR knockout pooled library (Addgene #73179), originally developed by David Root and John Doench, was used for amplification (Doench et al., 2016). A total of 100 ng of the library was introduced into 25 μL of Endura electrocompetent cells (Lucigen, #60242) in four separate transformations. Electroporation was carried out using a Biorad Micropulser at settings of 10 μF, 600 Ω and 1800 V. Immediately after electroporation, 975 μL of SOC medium was added within 10 seconds, after which the cells were incubated at 37 °C with shaking at 250 rpm for 1 hour. Following recovery, the cultures were plated on 245 mm square LB-agar dishes (Corning) supplemented with 100 μg/mL ampicillin and incubated at 32 °C for 14 hours. Colonies were harvested by rinsing the plates twice with 20 mL LB medium and scraping with a sterile cell scraper (Goh et al., 2021). The pooled cells were then processed for plasmid DNA extraction using the Macherey-Nagel Endotoxin-Free Plasmid Purification Kit (#740424-10).

### Mammalian cell culture

The A375 melanoma cell line (ATCC, CRL-1619) and HEK293FT cells (Invitrogen, R70007) were cultured every three days under standard conditions to sustain exponential growth. Cells were maintained in Dulbecco’s Modified Eagle Medium (DMEM; Gibco, 61965059) supplemented with 10% fetal bovine serum (FBS; Gibco, 10500064) and 1% penicillin-streptomycin (Merck, P4458), incubated at 37 °C with 5% CO_2_ in a humidified atmosphere.

### Lentiviral production

For lentivirus generation, HEK293FT cells were plated in 100 mm dishes to achieve approximately 70-80% confluency after 24 hours. Cells were cultured in DMEM as described above, with the addition of 2 mM L-glutamine (Gibco, 25030149) and 1 mM sodium pyruvate (Gibco, 11360070). Transfections were carried out using Lipofectamine 3000 reagent (Invitrogen, L3000001). The transfection cocktail consisted of 2 μg each of packaging plasmids PLP1 and PLP2, 1 μg of pVSVG envelope plasmid and 2 μg of the Brunello sgRNA library plasmid. These plasmids were mixed in 612.5 μL Opti-MEM (Gibco, 31985062) with 14 μL of Lipofectamine 3000 reagent and incubated at room temperature for 15 minutes prior to being added dropwise to the cells. Following transfection, the cells were returned to the incubator (37 °C, 5% CO_2_) for 24 hours. To enhance lentiviral production, the medium was supplemented with 1 mM sodium butyrate (Merck, 156-54-7) 24 hours post-transfection. Supernatants containing virus were collected at 48 and 72 hours, by centrifugation at 16,500 rpm for 90 minutes at 4 °C and the viral pellets were resuspended in PBS (Gibco, 20012027) before being stored at −80 °C. A375 cells were then used to determine the optimal multiplicity of infection (MOI) for downstream screening. Serial dilutions of the viral stock (1:10, 1:50, 1:100, and 1:1000) were prepared and added to 1×10^6^ A375 cells in 12-well plates along with 5 μg/mL polybrene (Merck, TR-1003-G). Plates were centrifuged at 3000 rpm for 90 minutes at 30 °C to enhance transduction efficiency. After 24 hours, the cells were split into two groups and cultured with or without 1 μg/mL puromycin. Following 72 hours of selection, viable cell counts were used to calculate transduction efficiency as followed:

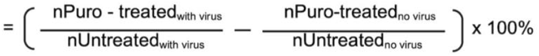

An MOI of 0.4 was selected for use in the large-scale screen.

### Cell viability assay

IC_50_ of Ganoderic Acid DM (Medchem, HY-120140) in A375 cells was evaluated using the PrestoBlue cell viability assay (Invitrogen, P50201), following the manufacturer’s instructions. A375 cells were plated in black 96-well plates at a density of 2,000 cells per well and incubated overnight to allow for cell attachment. Cells were then treated with either the vehicle control (Dimethyl sulfoxide (DMSO)) or with varying concentrations of Ganoderic Acid DM ranging from 1.6 μM to 100 μM and incubated for 24 or 48 hours. Cell viability was quantified by measuring metabolic activity via the PrestoBlue reagent, which generates a fluorescence signal corresponding to the number of viable cells (Lall et al., 2013). Fluorescence intensity was recorded at 560 nm using a microplate reader. All experiments were conducted in triplicates.

### GA-DM CRISPR/Cas9 screen

A total of 3.4×10^8^ A375 cells were transduced with the Brunello CRISPR knockout library at an MOI of 0.4 to achieve >400x library coverage. Cells were plated at 1×10^6^ per well across multiple 12-well plates for uniform infection, with each well receiving 12 µL of concentrated Brunello lentiviral preparation. After a 24-hour incubation, all wells were pooled into T175 flasks to maintain homogeneous library representation, with 1 µg/mL puromycin to select for successfully transduced cells. Selection was maintained for seven days. On day 7, cells were split into two experimental groups: 1) control cells treated with DMSO and 2) cells treated with 50 µM GA-DM. Each experimental group was prepared in duplicates with 3×10^7^ cells per replicate, maintaining >300x coverage. Cells were passaged every 3-4 days with freshly added DMSO or GA-DM. Finally, cell pellets were collected at both day 7 and day 14 post-treatment for downstream analysis (Goh et al., 2021).

### Genomic DNA extraction and amplification

Frozen A375 cell pellets previously stored at −80 °C were thawed, of which the genomic DNA (gDNA) was isolated using the Blood and Cell Culture DNA Maxi Kit (Qiagen, 13323), following the respective manufacturer protocols. PCR amplification was carried out using Q5 Hot Start High-Fidelity 2× Master Mix (NEB, M0494L), with 1.5 μg of gDNA input per reaction set.

For each sample, ten parallel 50 μL PCRs were prepared, each incorporating a unique forward primer containing a 1-10 bp staggered sequence to improve sequencing diversity. A single reverse primer containing a unique barcode was used to distinguish between samples (primer details listed in Table S1). Thermal cycling conditions included: an initial denaturation at 98 °C for 3 minutes, followed by 28 cycles of 98 °C for 10 seconds, 60 °C for 30 seconds and 72 °C for 25 seconds, concluding with a final extension at 72 °C for 5 minutes. The optimal number of cycles was determined in advance using gradient PCR to avoid amplification bias.

The ten PCRs per sample were combined and purified using the QIAquick PCR Purification Kit (Qiagen, 28104). Products were run on a 2% agarose gel and the target band (∼275 bp) was excised and purified using the Qiagen Gel Extraction Kit (Qiagen, 28704). Purified libraries were stored at −20 °C until they were submitted for next-generation sequencing, which was performed on the HiSeq-SE150 platform by NovogeneAIT Genomics (Singapore).

### Bioinformatics

Next-generation sequencing data in FASTQ (fastq.gz) format were demultiplexed by NovogeneAIT Genomics (Singapore). Sequence quality was assessed using FastQC, which provided insights into read quality, GC distribution and the presence of adapter sequences(Andrews et al., 2012). Following quality control, the FASTQ files were uploaded to the CRISPRAnalyzer web tool (http://crispr-analyzer.dkfz.de) for analysis (Winter et al., 2017). Within this platform, sgRNA reads were aligned to the Brunello reference library and corresponding read count files were generated to reflect sgRNA abundance. These counts were used to calculate log_2_ fold changes between treatment and control groups. sgRNAs with fewer than 20 reads were excluded to minimize noise in the dataset. To identify genes significantly affected by the treatment, the MAGeCK algorithm (Model-based Analysis of Genome-wide CRISPR/Cas9 Knockout) was employed (W. Li et al., 2014). MAGeCK generated gene rankings based on sgRNA enrichment or depletion, applying the Benjamini-Hochberg method to adjust p-values and control the False Discovery Rate (FDR). All analyses were conducted using default settings and genes with an FDR-adjusted p-value below 0.05 were considered statistically significant. To understand the biological functions of the identified genes, Gene Ontology (GO) enrichment analysis was performed (https://geneontology.org) and protein interaction networks were constructed using the STRING database, version 11.5 (https://string-db.org) (Aleksander et al., 2023; Ashburner et al., 2000; Szklarczyk et al., 2023).

### Data visualisation

Graphs and quantitative data were generated using RStudio (Posit, https://posit.co/products/open-source/rstudio/) and GraphPad Prism version 10.4.1 (GraphPad Software, https://www.graphpad.com). The chemical structure of GA-DM was drawn using ChemDraw software (https://revvitysignals.com/products/research/chemdraw). A schematic illustrating the potential mode(s) of action of GA-DM was created using BioRender (https://www.biorender.com).

## Results

To investigate the molecular mechanisms underlying the cytotoxic effects of GA-DM, we performed a genome-wide CRISPR/Cas9 knockout screen in A375 melanoma cells using the Brunello library. Prior to the screen, we determined the inhibitory concentration of GA-DM that reduces cell viability by 50% (IC_50_). As shown in figure 1, treatment with increasing concentrations of GA-DM for 48 hours resulted in a dose-dependent reduction in cell viability. A concentration of 50 µM GA-DM caused approximately 50% growth inhibition and was therefore selected for downstream experiments.

**Figure 1.**
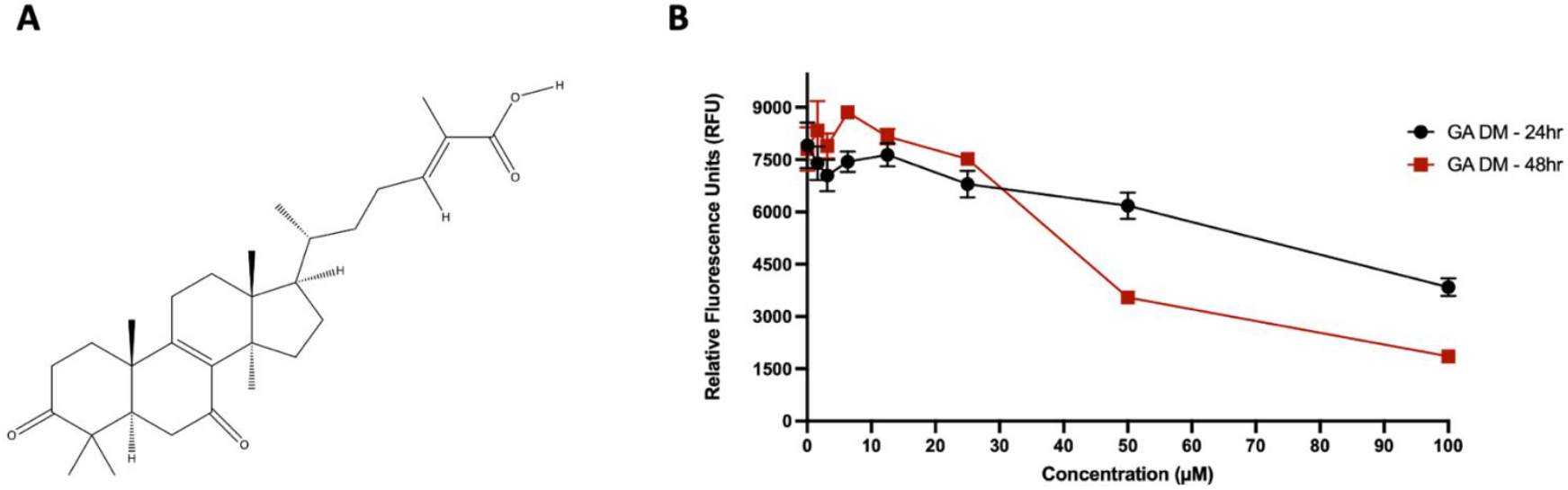
GA-DM’s effect on cell viability. A) Chemical structure of Ganoderic Acid DM. B) Dose-dependent effect of GA-DM on cell viability measured by PrestoBlue assay at 24 and 48 hours. Cells were treated with increasing concentrations of GA-DM (0-100 µM) and relative fluorescence units (RFU) were measured as a readout of metabolic activity. Data represent mean and SD of biological replicates. A time-and dose-dependent decrease in viability is observed, indicating GA-DM induced cytotoxicity.

The Brunello library contains 76,441 sgRNAs targeting 19,114 genes, along with 1,000 non-targeting controls (Doench et al., 2016). A375 cells were transduced with the lentiviral library at a multiplicity of infection (MOI) of ∼0.4 to achieve the recommended representation of ∼400 cells per sgRNAs. After puromycin selection for 7 days to eliminate uninfected cells, populations were treated with either DMSO (control) or 50 µM GA-DM in biological duplicates and samples were collected after 7 and 14 days of treatment. Genomic DNA was extracted, sgRNA sequences were PCR-amplified and next-generation sequencing (NGS) was performed to assess changes in sgRNA abundance. Enriched sgRNAs represent genes whose loss conferred resistance to GA-DM, while depleted sgRNAs indicate genes whose knockout sensitised cells to the compound.

NGS data were analysed using the CRISPRAnalyzeR platform, with the MAGeCK algorithm used to identify statistically significant sgRNA enrichments or depletions (fig. 3) (Li et al., 2014; Winter et al., 2017). Read count distributions across all samples showed comparable representation and sequencing depth, confirming uniform library coverage (fig. S1 and S2). The day 7 duplicates demonstrated strong reproducibility across the screen (fig. S3A). The DMSO-treated samples showed a Pearson correlation of 1 and a Spearman correlation of 0.97-0.98, while the GA-DM 50 µM-treated samples exhibited similarly high reproducibility, with a Pearson value of 1 and a Spearman value of 0.96. Likewise, the day 14 duplicates from the resistance screen (fig. S3B) also showed strong pairwise correlations. The DMSO-treated samples maintained a Pearson correlation of 1 and a Spearman correlation of 0.97, while the GA-DM 50 µM-treated samples displayed a Pearson value of 1 and a Spearman value of 0.93.

**Figure 2.**
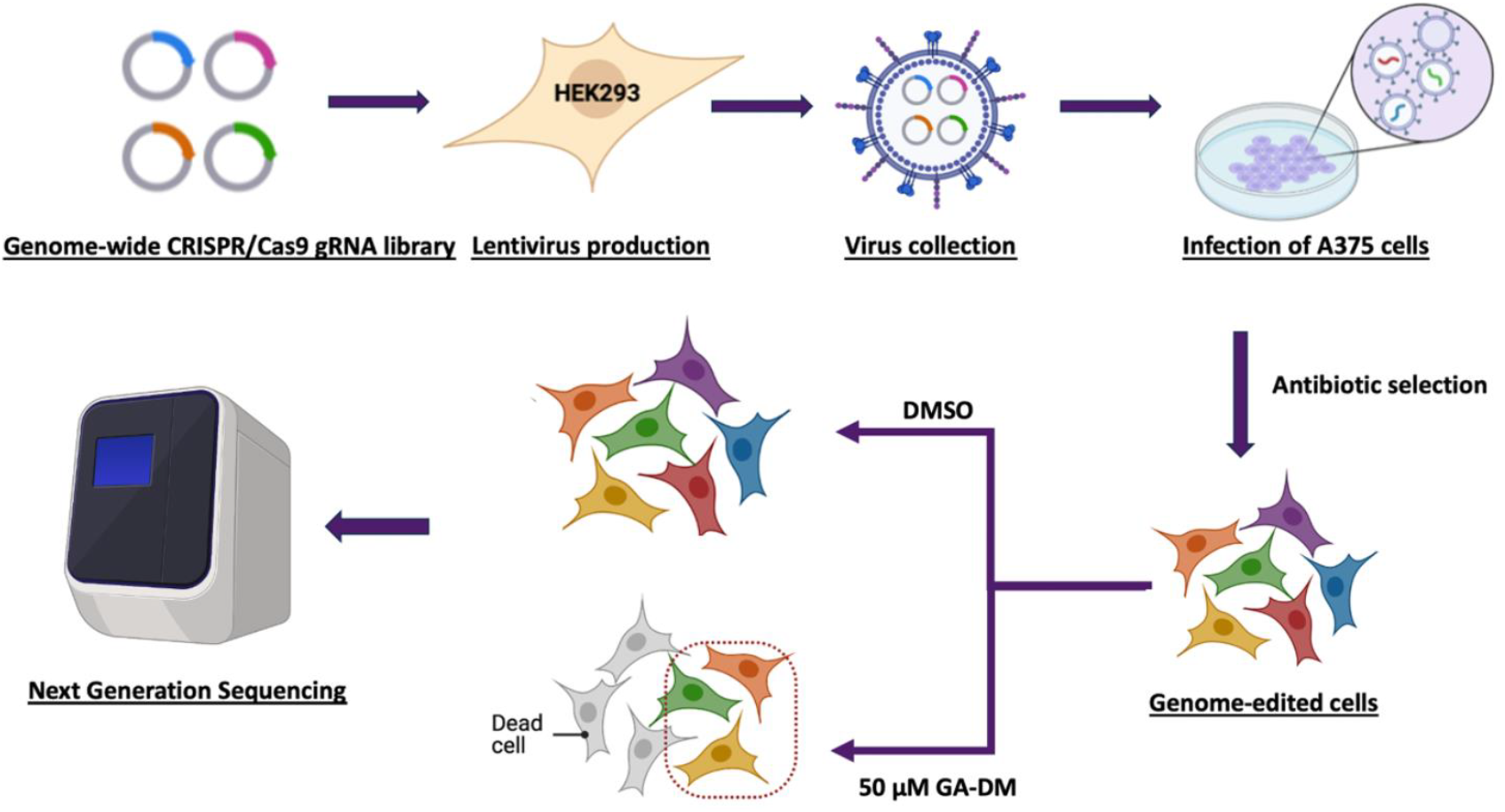
Schematic overview of the genome-wide CRISPR/Cas9 screen workflow. Briefly, A375 cells were transduced with the Brunello human CRISPR knockout pooled lentiviral library, which contains 76,441 sgRNAs targeting 19,114 genes. Following puromycin selection, the cells were treated with DMSO or 50 µM GA-DM and collected at day 7 and day 14, with each group having two biological replicates. Finally, genomic DNA samples were harvested from the initial infected cell population and from those treated with DMSO or GA-DM, amplified by PCR and deep-sequenced by next-generation sequencing (NGS).

**Figure 3.**
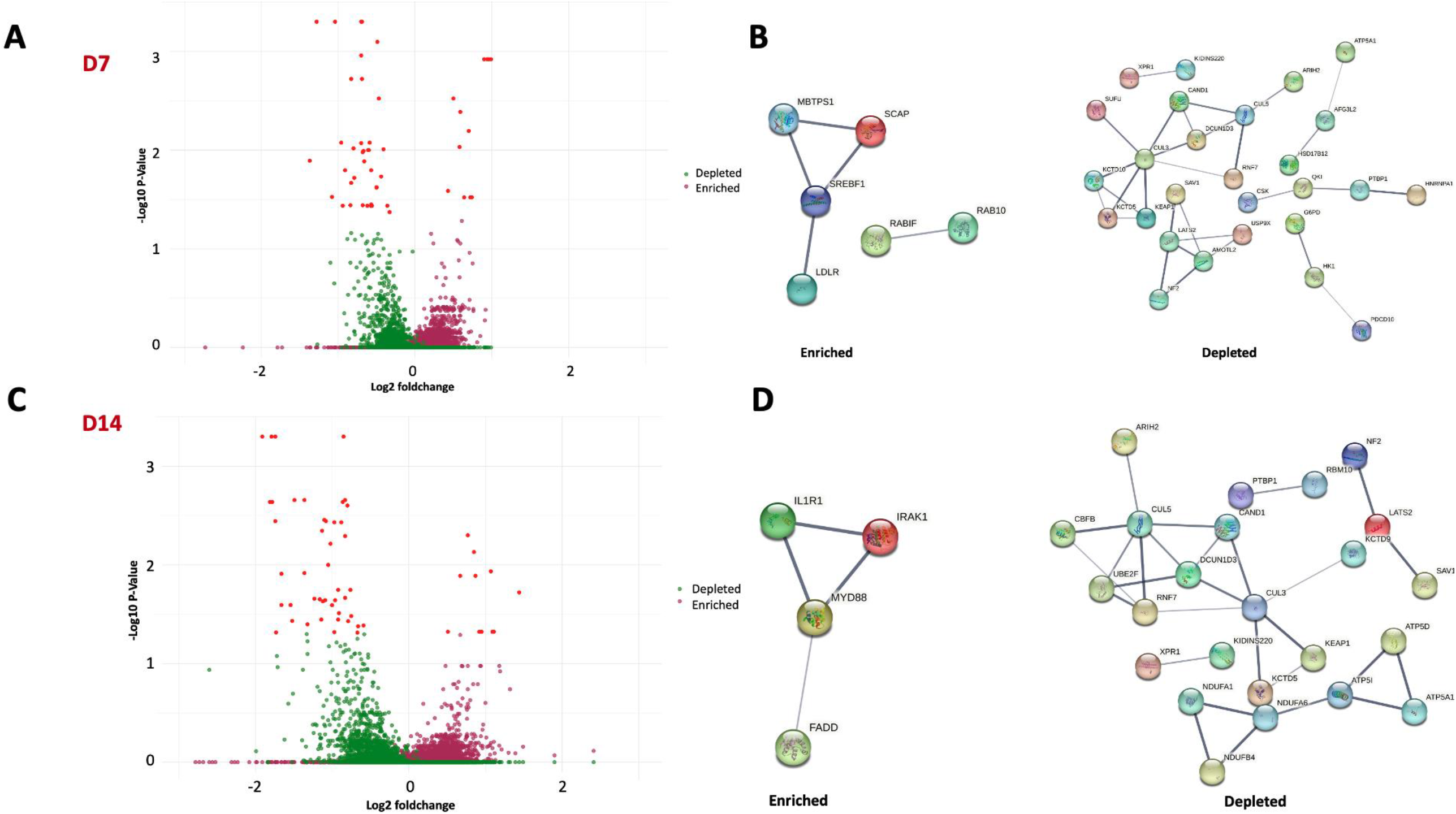
Genome-wide CRISPR screen identifies genes modulating sensitivity to GA-DM at day 7 and day 14. The sgRNA read count distribution of DMSO-treated and GA-DM-treated cells were analysed using MAGeCK. A) Volcano plot displaying the CRISPR screen results of day 7. The x-axis represents log_2_ fold change and the y-axis indicates -log_10_ *p* value. Green dots represent depleted genes, while purple dots indicate the enriched genes. Significant genes (*p* <0.05) are highlighted in red. B) STRING network analysis of the top significantly enriched (left) and depleted (right) genes at D7, revealing functionally connected pathways. C) Volcano plot of the CRISPR screen results at day 14. Significant genes (*p* <0.05) are again marked in red, with green and purple indicating depleted and enriched genes, respectively. D) STRING network of significantly enriched and depleted genes of day 14.

After 7 days of GA-DM treatment, 12 genes showed significantly enriched sgRNAs (p < 0.05), while 63 genes had significantly depleted sgRNAs (p < 0.05) (fig. 3A, table S2, S3). At day 14, 13 genes were significantly enriched and 71 genes were significantly depleted (fig. 3D, table S7, S8). To explore potential functional interactions, the top-ranked enriched and depleted genes (based on p-value and log_2_ fold change) were analysed using STRING v11.5 (Szklarczyk et al., 2023). The 7-day resistance hits formed a significantly interconnected protein-protein interaction (PPI) network (PPI enrichment p = 4.99×10^4^). Gene Ontology (GO) enrichment analysis revealed that these genes were primarily involved in the SREBP signalling pathway (e.g., SREBF1, SCAP) and regulation of cholesterol metabolism (*MBTPS1, LDLR, SCAP* and *SREBF1)* (fig. 3C, table S4). Similarly, the 63 significantly depleted genes from the 7-day GA-DM treatment group also displayed a strong PPI enrichment (p = 7.64×10^-9^), indicating functional relatedness (fig. 3C, table S5, S6). These genes were associated with the negative regulation of LDL receptor activity (*CSK, MYLIP*), the Hippo signalling pathway (*SAV1, NF2* and *LATS2)* and ubiquitin mediated proteolysis (*KEAP1, CUL3, RNF7, FBXW7, TRIP12* and *CUL5*). At the 14-day timepoint, enriched genes showed a PPI enrichment value of 8.59×10^-5^ and were primarily linked to inflammatory and immune signalling pathways, including NF-κB signalling (*NFKB1, IRAK1, IL1R1, MYD88)* and the Toll-like receptor pathway (*NFKB1, IRAK1, FADD* and *MYD88)* (fig. 3D, table S9). The 71 significantly depleted genes had a PPI enrichment value of 1.63×10^-8^ and included genes involved in protein neddylation (*RNF7, DCUN1D3, UBE2F*) and mitochondrial ATP synthesis (*ATP5D, ATP5I, ATP5A1*) (table S10, 11). Notably, components of the Hippo pathway (*SAV1, NF2, LATS2*) remained significantly depleted at both time points, suggesting a consistent role in mediating sensitivity to GA-DM.

Although downstream experimental validation was not performed in this study, the integration of CRISPR screen hits with functional annotation and known signalling pathways provides valuable insights into the potential mechanisms of action of GA-DM. These findings lay the groundwork for further investigation into the roles of these candidate genes and pathways in modulating GA-DM activity.

## Discussion

Ganoderic acid DM (GA-DM), a triterpenoid derivative of *Ganoderma lucidum*, has been studied for its diverse therapeutic effects, particularly its anticancer, anti-inflammatory and metabolic properties. However, the molecular mechanisms underlying its cytotoxic activity remain poorly defined. In this study, we have found that genes involved in regulating the lipid homeostasis (particularly, the SREBP pathway) and the inflammatory pathway modulate cellular sensitivity to GA-DM. In addition, genes encoding components of the ubiquitination machinery and Hippo signalling were found to significantly influence GA-DM sensitivity.

Strikingly, several genes involved in cholesterol biosynthesis and regulation (*SREBF1, SCAP, MBTPS1, LDLR)* emerged as resistance hits. These genes are central to the SREBP pathway which regulates cholesterol and fatty acid homeostasis (Silvente-Poirot & Poirot, 2012). Under low cholesterol conditions, SREBP gets activated and promotes expression of target genes involved in lipid biosynthesis (fig. 4) (Brown & Goldstein, 1997). Knockout of these genes likely reduces cholesterol synthesis, suggesting that GA-DM may rely on intact lipid metabolic pathways to exert cytotoxic effects. In this context, loss of cholesterol biosynthesis may mitigate GA-DM-induced cellular stress, thereby conferring resistance. These findings led us to propose a novel mechanism of action for GA-DM, in which it disrupts cellular lipid homeostasis to cause cytotoxicity. This aligns with prior observations that GA derivatives can influence lipid metabolism, including reduced cholesterol levels and inhibition of key metabolic enzymes (Zhu et al., 2018). While previous work has largely emphasised GA’s effects on apoptosis, inflammation and immune modulation, our results provide the first genome-scale functional evidence linking cholesterol metabolism to GA-DM sensitivity.

**Figure 4.**
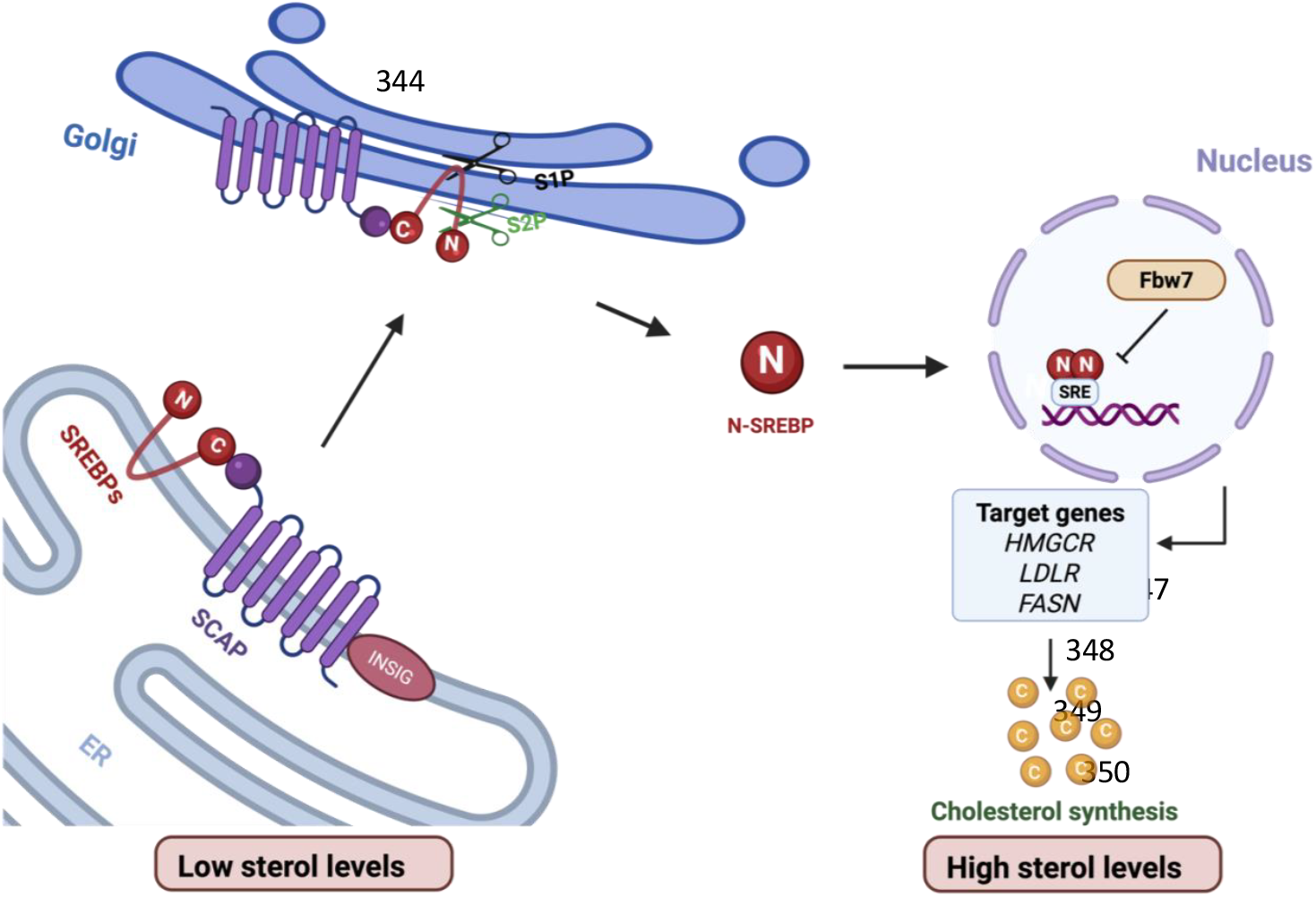
The SREBP-cholesterol regulatory pathway. Under low sterol conditions, SREBPs (Sterol Regulatory Element-Binding Proteins) form a complex with SCAP in the endoplasmic reticulum (ER), which facilitates their transport to the Golgi apparatus. There, SREBPs undergo sequential proteolytic cleavage by Site-1 (S1P) and Site-2 (S2P) proteases, releasing the active N-terminal fragment (N-SREBP). This fragment translocates to the nucleus, where it binds sterol regulatory elements (SREs) and activates the transcription of genes involved in cholesterol and lipid biosynthesis, including HMGCR, LDLR, and FASN, ultimately increasing cellular cholesterol levels.

Further supporting this mechanism, the screen also identified sensitivity hits in genes involved in ubiquitin-mediated proteolysis, including *FBXW7* and *MYLIP*, both of which are known to regulate lipid metabolism through SREBP turnover. For instance, *FBXW7* targets phosphorylated nuclear SREBPs for degradation and its loss has been shown to stabilise SREBPs and promote lipid accumulation (Bengoechea-Alonso et al., 2022; Sundqvist et al., 2005). Similarly, *MYLIP* (also known as *IDOL*) ubiquitinates the LDL receptor, limiting cholesterol uptake (Zelcer et al., 2009). Their depletion in our screen sensitises cells to GA-DM, consistent with the hypothesis that enhanced cholesterol synthesis or uptake may enhance GA-DM toxicity. Furthermore, components of the Hippo signalling pathway (*LATS2, SAV1, NF2*) were also found to confer sensitivity to GA-DM. This pathway, known for regulating organ size, cell proliferation and apoptosis, has also been implicated in lipid metabolism (Udan et al., 2003). Specifically, *LATS2* has been shown to inhibit SREBP activation by binding to and retaining it in an inactive state (Aylon et al., 2016). Therefore, loss of *LATS2* likely results in increased SREBP activity and lipid synthesis, which could exacerbate GA-DM-induced stress and explain the observed sensitisation upon gene knockout.

Another key finding from our CRISPR screen was the enrichment of genes involved in inflammatory signalling, particularly components of the NF-κB and Toll-like receptor (TLR) pathways, as resistance hits. This implies that GA-DM may engage or activate inflammatory responses as part of its cytotoxic mechanism. Of interest, these hits were predominantly enriched at later time points (day 14), suggesting a temporal progression in which initial metabolic disruption by GA-DM leads to a delayed inflammatory response. Emerging evidence supports significant crosstalk between SREBP-driven lipid metabolism and inflammation, particularly through NF-κB signalling (Fei et al., 2023; Ferré & Foufelle, 2007; He et al., 2017; Oishi et al., 2017; Reboldi et al., 2014). NF-κB activation has been shown to promote intracellular cholesterol accumulation and enhance LDL uptake, while cholesterol overload itself can amplify pro-inflammatory signalling, indicating a potential feed-forward loop between these pathways (He et al., 2017; Reboldi et al., 2014). However, this relationship appears to be highly context-dependent, as NF-κB activation has been associated with both upregulation and suppression of SREBP activity depending on cell type and conditions (Fei et al., 2023). Moreover, TLRs are known to activate NF-κB via both MyD88-dependent and independent signalling cascades (Kawasaki & Kawai, 2014). Intriguingly, MyD88 signalling has also been implicated in promoting cholesterol biosynthesis through activation of the AKT/mTOR–SREBP axis, suggesting a molecular bridge between innate immune responses and lipid metabolism (Hsieh et al., 2020; Zhou et al., 2020). Although the details of this mechanistic interplay have not been fully elucidated, our findings support a model in which inflammatory signalling, particularly through TLRs and NF-κB, not only contributes to but may also potentiate lipid dysregulation in response to GA-DM. This interplay between metabolic and inflammatory stress could be a critical component of GA-DM’s cytotoxic activity.

We propose a set of working models in which GA-DM cytotoxicity is dependent on cholesterol or specific lipid environments (fig. 5). This is supported by our findings that cellular sensitivity to GA-DM is modulated by disruptions in cholesterol and lipid metabolism, whether through inhibition of biosynthetic enzymes, altered protein turnover or altered inflammatory signalling. In model I, GA-DM interferes with the processing of SREBPs under low sterol conditions, thereby blocking SREBP cleavage and nuclear translocation. This prevents the transcriptional activation of lipid and sterol biosynthesis genes, ultimately disrupting lipid homeostasis. In this model, treatment of SREBP1 knockout cells with GA-DM will have no further inhibitory effect resulting in ‘resistance’ to GA-DM. In model II, GA-DM instead promotes SREBP activation, resulting in excessive upregulation of lipid and cholesterol biosynthesis to toxic levels. Deletion of SREBP1 will hence lead to resistance to GA-DM. These opposing models suggest that GA-DM may either suppress or hyperactivate the SREBP pathway depending on cellular context. Experimental analysis of SREBP activation status under different sterol conditions in the presence of GA-DM will be essential to distinguish between these possibilities. Finally, model III posits a distinct but complementary mechanism, whereby GA-DM preferentially associates with cholesterol- or lipid-rich membranes. In this scenario, lowering intracellular lipid or cholesterol content disrupts GA-DM’s membrane binding and confers resistance. This model yields two testable predictions: (1) inhibition of cholesterol biosynthesis should increase resistance to GA-DM and (2) GA-DM should exhibit selective binding to cholesterol-rich membranes. These predictions can be experimentally tested by determining the effect of lovastatin treatment on resistance to GA-DM and *in vitro* liposome-binding assays with defined lipid compositions, respectively. Together, these models provide a framework for dissecting the cholesterol-dependent mechanisms that underlie GA-DM-induced cytotoxicity and the basis for their anti-obesity therapeutic properties.

**Figure 5.**
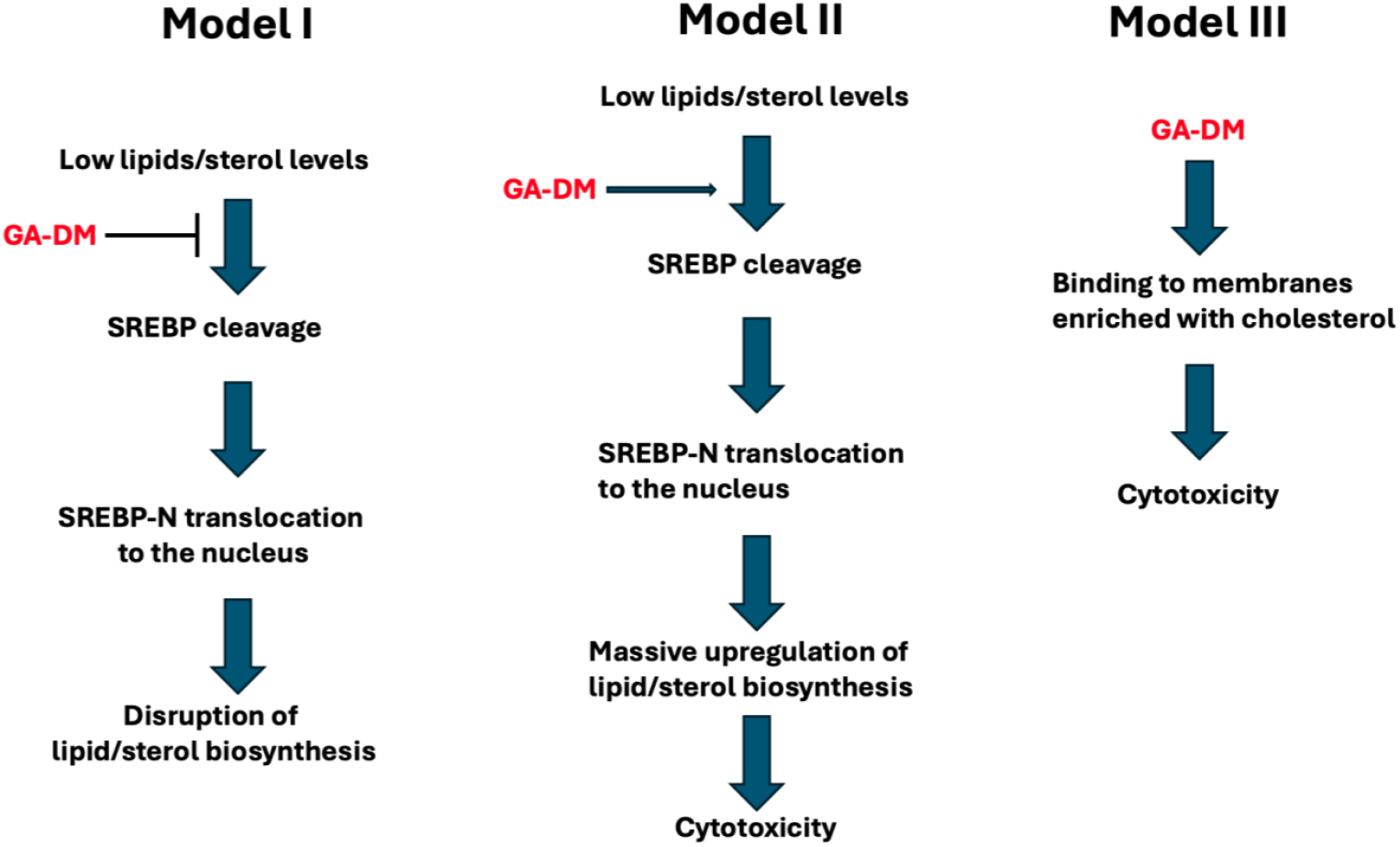
Proposed working models for GA-DM-induced cytotoxicity involving cholesterol and lipid metabolism. We present three models to explain the cholesterol-dependent cytotoxic effects of GA-DM. In model I, GA-DM inhibits SREBP processing under low sterol conditions, preventing nuclear translocation and transcriptional activation of lipid and sterol biosynthesis genes, thereby disrupting lipid homeostasis. In Model II, GA-DM enhances SREBP activation, leading to toxic lipid and cholesterol accumulation. These opposing models suggest that GA-DM may either suppress or enhance the SREBP pathway depending on context. In Model III, GA-DM preferentially binds to cholesterol- or lipid-rich membranes; lowering intracellular lipid or cholesterol levels may disrupt this interaction and confer resistance. This model predicts that GA-DM binding can be modulated by cholesterol biosynthesis inhibitors (e.g. lovastatin) and can be tested using *in vitro* liposome-binding assays.

In summary, our genome-wide CRISPR screen identified a link between GA-DM and pathways regulating cholesterol biosynthesis and inflammatory signalling in human cells. Although these findings offer valuable insights, additional mechanistic studies are required to elucidate the specific molecular pathways underlying these associations.

## Conclusion

This study identifies novel genetic determinants of cellular resistance and sensitivity to GA-DM, implicating lipid metabolism, protein ubiquitination and inflammatory signalling as key modulators of its action. The integration of unbiased CRISPR screening with pathway-level analyses provides new insight into the possible biological mechanisms of GA. Future work should focus on validating these findings through targeted gene knockouts, cholesterol/lipid assays and pharmacological studies, potentially paving the way for therapeutic combinations (e.g. with statins or proteasome inhibitors) that could enhance GA-DM efficacy. More broadly, these results highlight the value of combining traditional compounds with modern genomic tools to uncover mechanistic insights that inform therapeutic development.

## Data availability statement

Strains and plasmids are available upon request. Supplemental information is provided, including genome-wide CRISPR/Cas9 oligos, lists of enriched and depleted genes at multiple time points, and functional enrichment analyses (GO terms and KEGG pathways). The raw sequencing reads from the CRISPR screen will be deposited in a public repository (e.g., NCBI Sequence Read Archive, ENA, or DDBJ) upon acceptance of the manuscript. Accession numbers will be provided in the final published version. The authors affirm that all other data necessary for confirming the conclusions of the article are present within the article, figures, and tables.

## Acknowledgements

NA was supported by a studentship jointly funded by University of Warwick and A*STAR. This project was funded by a SIFBI seed grant awarded to JL and PA. The authors are grateful to Dr Manikandan Lakshmanan (IMCB) for providing access to his tissue culture lab facilities for the project.

## Author’s contribution

NA: project conceptualisation, methodology, investigation, data analysis and writing (original draft, review and editing). JL: project conceptualisation, methodology and investigation. PA: project conceptualisation, funding acquisition, methodology, supervision and writing (review and editing). All authors reviewed the manuscript.

## Supplementary Information

Supplementary tables are provided in the accompanying excel files.

### Figures

**Figure S1.**
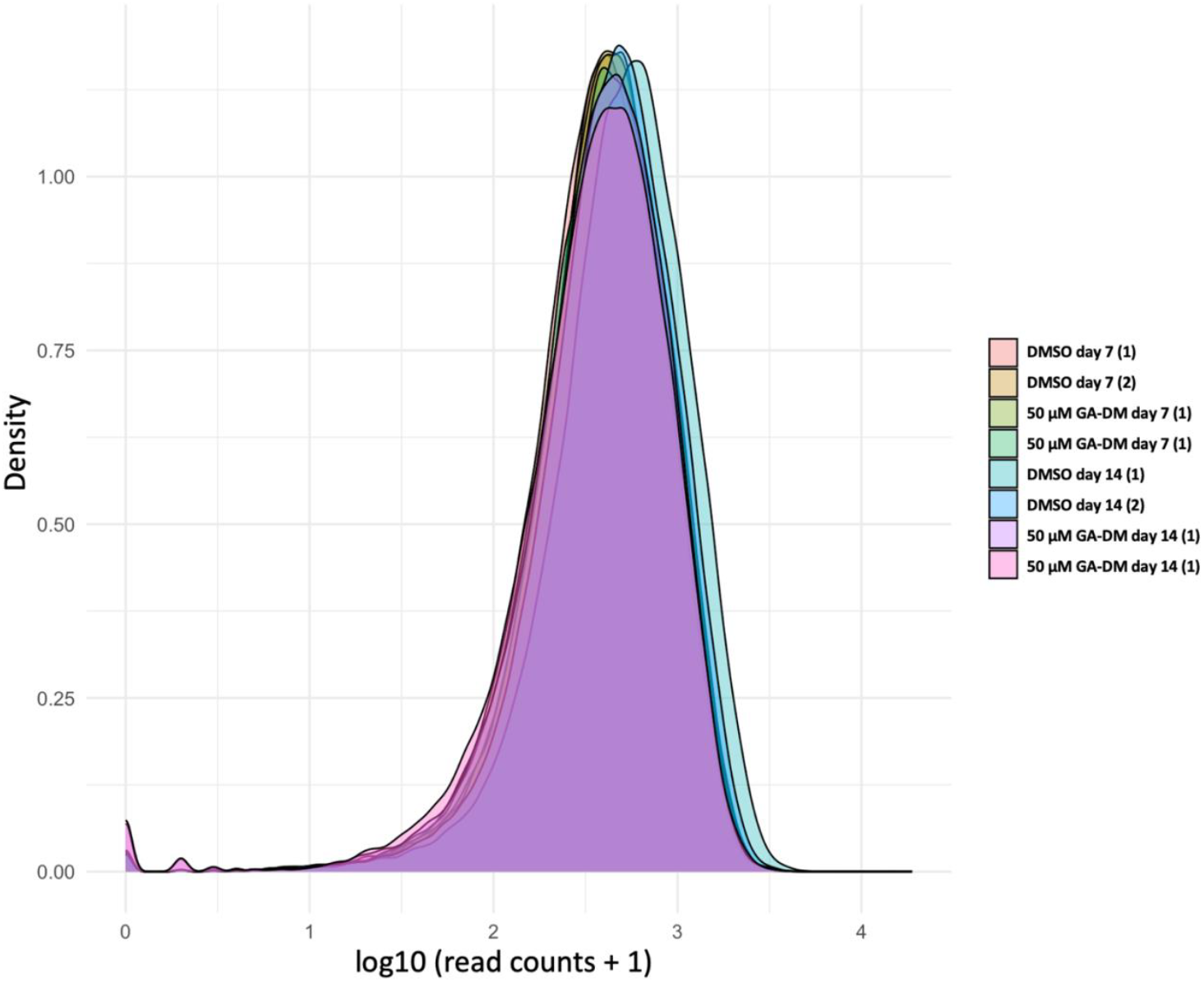
Distribution of sgRNA read counts across CRISPR screen samples. Density plots showing the distribution of log_10_-transformed sgRNA read counts (log_10_[read counts + 1]) for all samples from the genome-wide CRISPR/Cas9 screens. Each curve represents one biological replicate from cells treated with DMSO or 50 µM GA-DM at day 7 or day 14. All samples exhibited similar unimodal distributions, indicating consistent library representation and sequencing depth across replicates and treatment conditions.

**Figure S2.**
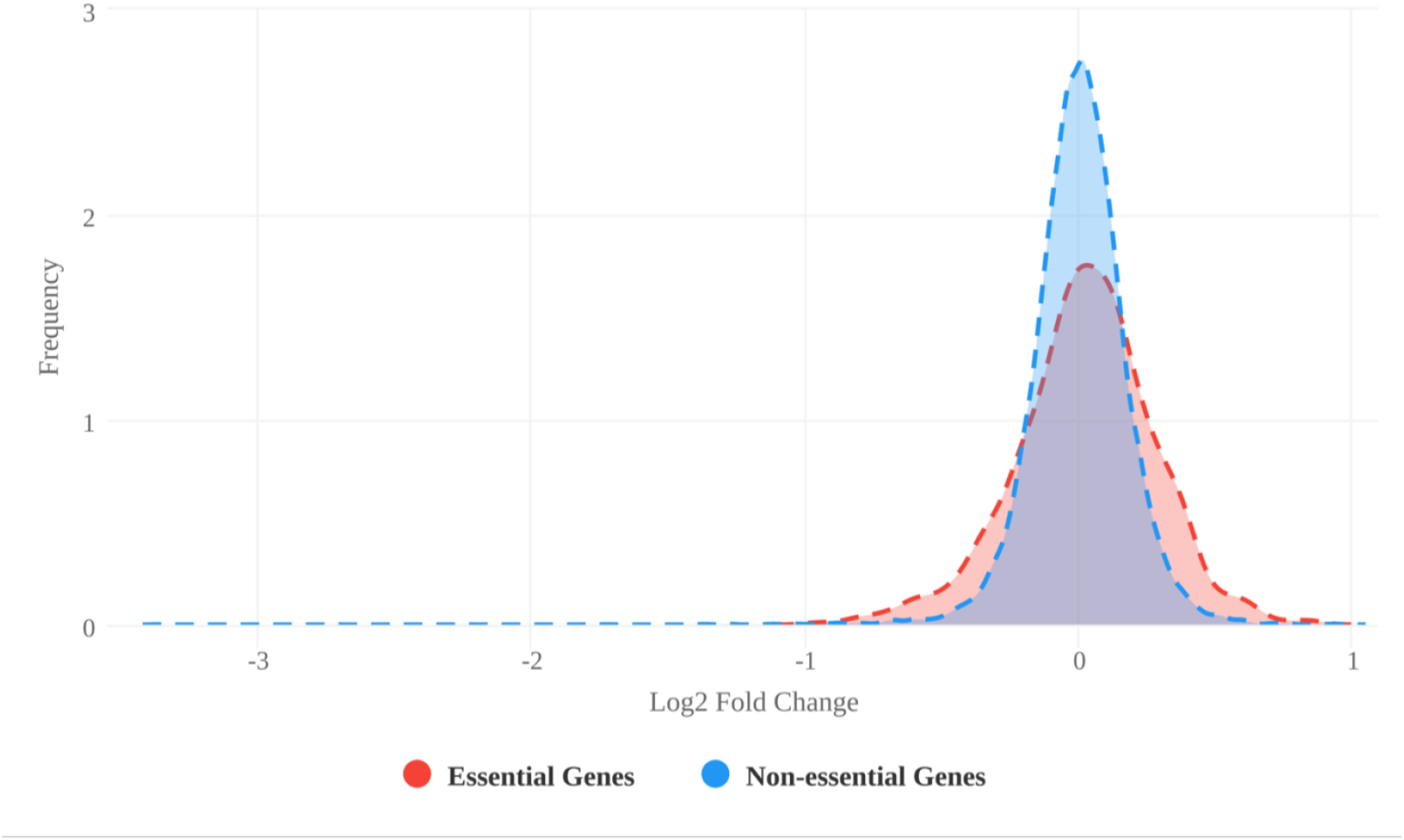
Distribution of Log_2_ Fold Changes for Essential and Non-essential Genes Identified by MAGeCK Analysis. Kernel density estimation plots showing the distribution of log_2_ fold changes for essential (red) and non-essential (blue) genes across the CRISPR screen, as determined by MAGeCK analysis. Essential genes display a broader distribution with greater negative fold changes, reflecting sgRNA depletion due to loss of cell viability, while non-essential genes are centred around zero, indicating minimal effect on cell survival. These distributions confirm the expected behaviour of control gene sets and validate the quality of the CRISPR screen data.

**Figure S3.**
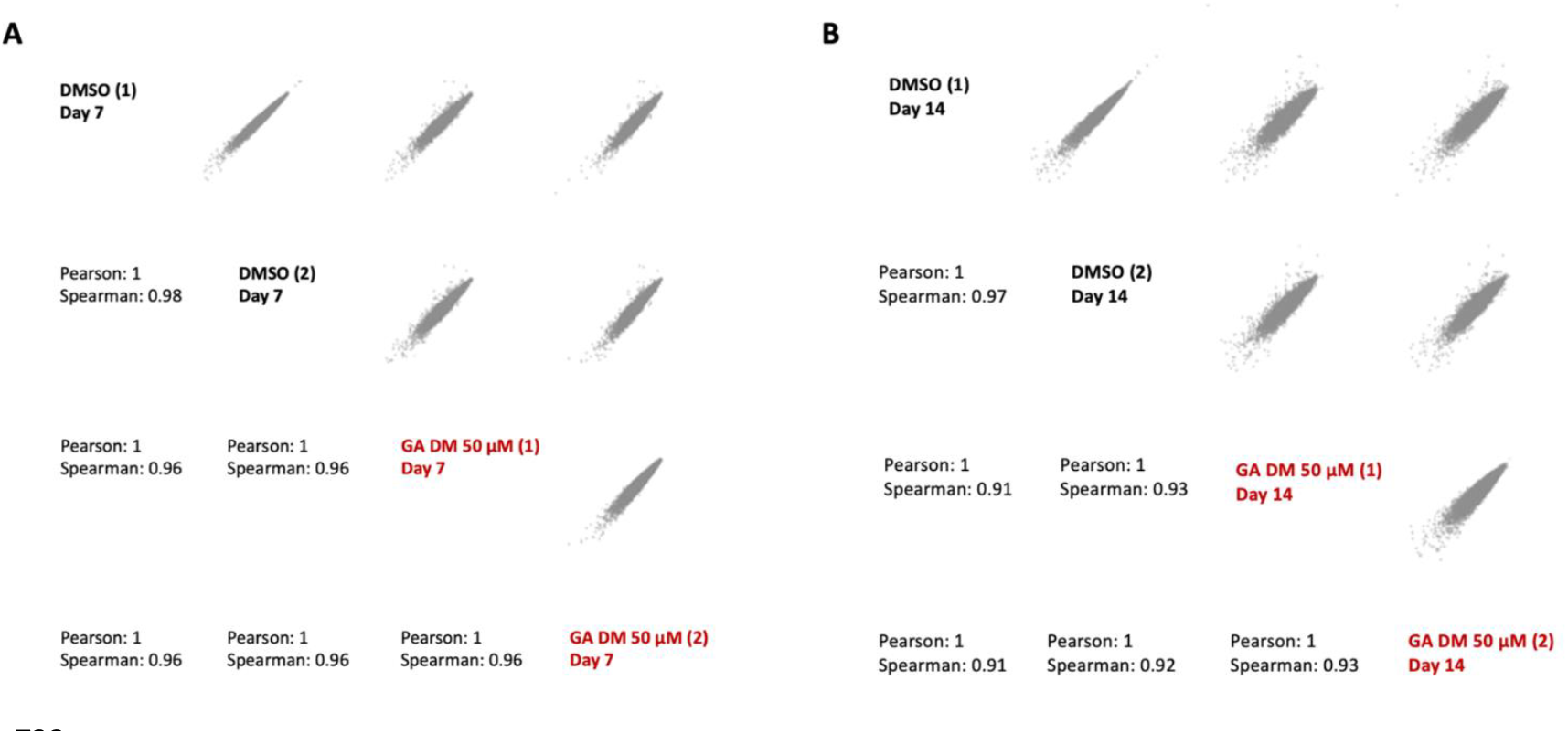
Pairwise correlations of CRISPR screen replicates. (A) Day 7 and (B) Day 14 replicate comparisons showing Pearson and Spearman correlation coefficients for DMSO- and 50 µM GA-DM-treated samples. Both conditions exhibited strong correlations across replicates, confirming high reproducibility and consistent cellular responses in the CRISPR screens.

